# Single-cell transcriptomics reveal cell type-specific molecular changes and altered intercellular communications in chronic obstructive pulmonary disease

**DOI:** 10.1101/2021.02.23.432590

**Authors:** Qiqing Huang, Jingshen Wang, Shaoran Shen, Yuanyuan Wang, Yan Chen, Shuangshuang Wu, Wei Xu, Bo Chen, Mingyan Lin, Jianqing Wu

**Affiliations:** Key Laboratory of Geriatrics of Jiangsu Province, Department of Geriatrics, the First Affiliated Hospital of Nanjing Medical University, Nanjing, Jiangsu, 210029, China; Department of Neurobiology, School of Basic Medical Sciences, Nanjing Medical University, Nanjing, Jiangsu 211166, China; Department of Respiratory and Critical Care Medicine, BenQ Medical Center, The Affiliated BenQ Hospital of Nanjing Medical University, Nanjing, Jiangsu Province, 210019, China

**Keywords:** chronic obstructive pulmonary disease (COPD), scRNA-seq, aging, smoking, monocyte, club cell, macrophage

## Abstract

Chronic obstructive pulmonary disease (COPD) is a common and heterogeneous respiratory disease, the molecular complexity of which remains poorly understood, as well as the mechanisms by which aging and smoking facilitate COPD development. Here, using single-cell RNA sequencing of more than 65,000 cells from COPD and age-stratified control lung tissues of donors with different smoking histories, we identified monocytes, club cells, and macrophages as the most disease-, aging-, and smoking-relevant cell types, respectively. Notably, we found these highly cell-type specific changes under different conditions converged on cellular dysfunction of the alveolar epithelium. Deeper investigations revealed that the alveolar epithelium damage could be attributed to the abnormally activated monocytes in COPD lungs, which could be amplified via exhaustion of club cell stemness as ages. Moreover, the enhanced intercellular communications in COPD lungs as well as the pro-inflammatory interaction between macrophages and endothelial cells indued by smoking could facilitate signaling between monocyte and the alveolar epithelium. Our findings complement the existing model of COPD pathogenesis by emphasizing the contributions of the previously less appreciated cell types, highlighting their candidacy as potential therapeutic targets for COPD.

## Introduction

Chronic obstructive pulmonary disease (COPD), characterized by persistent airflow limitation and incompletely reversible airway construction, is a common and heterogeneous respiratory disease (1). As a leading cause of chronic morbidity and mortality throughout the world, COPD represents a major public health problem and challenge (2, 3).

Although a large amount of work has been done, our understanding of the pathological mechanisms of COPD is still limited, which is partially attributed to the anatomical, histological, and physiological complexity of the respiratory system. More than 40 cell types are involved in the composition of lung tissue, performing their own duties as well as forming precise temporal and spatial interaction networks to maintain the homeostasis of lung (4, 5). Moreover, in addition to serving as a vital organ for gas exchange that is continually exposed to external environment, lung also functions as a physical and immune barrier against various exogenous pathogens via cooperating closely with immune system (6). As a widely accepted systemic inflammatory disorder, COPD is undoubtedly associated with chronic and abnormal inflammatory responses contributing to airway disease and emphysema. Hence, previous studies paid most attention to the role of macrophages and epithelial cells in COPD (7). Recently, Shibata et al. reported that basophils trigger emphysema development through IL-4-mediated generation of MMP-12-producing macrophages, implying that previously unappreciated cell type might play an important role in COPD development (8).

The prevalence of COPD rises dramatically with age (9) and tobacco use (10), making age and cigarette smoking high risk factors for COPD (11). Indeed, there are many clinical similarities between aged lungs and COPD lungs, including decline of lung function, decreased mucociliary clearance, ongoing inflammation, unbalanced oxidative stress, compromised vascular regulation, alveolar enlargement, and altered extracellular matrix (ECM) components (9, 12). COPD thus has long been proposed to be a disease of accelerated lung aging (9). Despite great efforts to prove this hypothesis, aging along is still insufficient to explain pathology of COPD, which in turn reminds us of the possibility that there may be some disease-specific factors mediating the contribution of aging to COPD development. Likewise, for the last decades, great steps have been taken to understand how smoking influences different kinds of cells in the lungs between individuals with or without COPD, but most of these studies focused on some well-known smoking-prone cell types (such as macrophages and T cells) or simple point-to-point crosstalk between these cell types (13), leaving the far more complex intercellular communications in reality that promote the development of COPD in smokers largely elusive.

In the present study, thanks to the progress in high-throughput techniques, in particular single-cell RNA Sequencing (scRNA-seq), we sought to elucidate the cell-type-specific molecular mechanisms underlying COPD at single-cell resolution, taking aging as well as smoking into account, via analyzing single-cell transcriptomic data of more than 65,000 cells in lung tissues from 3 patients with COPD, 3 normal elderly donors, and 3 normal young donors. We revealed the altered intercellular communications within COPD lungs and attribute apoptosis of alveolar epithelial cells responsible for the alveolar wall destruction to abnormal activation of monocytes, the former of which was enhanced by the exhausted regeneration and differentiation of club cells under the condition of lung aging and the enhanced interaction between macrophages and endothelial cells induced by smoking. Our study offers new insights into COPD pathogenesis and elucidates the mechanisms underlying the promotive effect of aging and smoking on COPD development in terms of cellular interaction at single-cell level.

## Methods

### Patients

This study was approved by the Ethics Committee at the First Affiliated Hospital of Nanjing Medical University (IRB-GL1-AF08). We complied with all relevant ethical regulations and written informed consent was obtained from each patient prior to surgery. Only patients with untreated, primary, non-metastatic lung tumors that underwent lung lobe resection with curative intent were included and were divided into three groups according to lung function and age: COPD (according to the GOLD guidelines) group (62 ± 11.53), normal elderly (≥ 65 years) group (73 ± 2.00), and normal young (≤ 40 years) group (30 ± 4.36). In order to get rid of the influence of cancer, lung tissues were isolated from more than 5 cm away from the tumor border. No patients had a history of asthma or renal dysfunction. The clinical characteristics of the patients are shown in Supplementary Table S1.

### Generation of single-cell suspensions from lung tissues

All samples were procured immediately upon surgery and were processed within 5 hours after procurement. Upon arrival to the laboratory, lung tissue (up to 20 × 20 × 20 mm) was washed with icy sterile PBS and was processed to prepare a single-cell suspension. Briefly, lung tissue was cut into pieces and incubated with 2 mg/ml collagenase I, 2 mg/ml collagenase IV and 0.2 mg/ml DNase I (Roche) in RPMI 1640 for 40 min at 37℃ with 150 rpm shaking. The resulting cell suspension was filtered through a 70-μm sterile nylon strainer and centrifuged at 400 g for 5 min at 4℃. The cell pellet was resuspended in 1 ml RBC lysis buffer (0.15 mol/l NH4Cl, 10 mmol/l KHCO3, 0.1 mmol/l EDTA) and lysed for 10 min at room temperature. Cells were centrifuged at 400 g for 5 min at 4℃. The cell pellet was resuspended in 400 μl PBS for cell sorting by Countstar (IC1000).

### Single-Cell RNA sequencing cell capture and cDNA synthesis

Using BD Rhapsody™ Cartridge Reagent Kit (BD, 633731) and BD Rhapsody™ Cartridge Kit (BD, 633733), the cell suspension (300-600 living cells per microliter determined by Count Star) was loaded onto the Rhapsody™ Cartridge (BD) to generate single-cell magnetic beads in the microwells according to the manufacturer’s protocol. In short, single cells were suspended in sample buffer (BD). About 18,000 cells were added to each channel, and the target cell will be recovered was estimated to be about 9,000 cells. Captured cells were lysed and the released RNA were barcoded through reverse transcription in individual microwells. Reverse transcription was performed on a ThermoMixer® C (Eppendorf) at 1,200 rpm and 37°C for 45 minutes. The cDNA was generated and then amplified, and quality assessed using an Agilent 4200 (performed by CapitalBio Technology, Beijing). Single-cell RNA sequencing (scRNA-seq) library preparation according to the manufacture’s introduction, scRNA-seq libraries were constructed using BD Rhapsody™ WTA Amplification Kit (BD, 633801). The libraries were finally sequenced using an Illumina Novaseq6000 sequencer with a sequencing depth of at least 50,000 reads per cell with pair-end 150 bp (PE150) reading strategy (performed by CapitalBio Technology, Beijing).

### Processing of scRNA-seq data

The BD Rhapsody analysis pipeline was used to process sequencing data (fastq files), the reference genome was GENCODE v29 (14). The scRNA-seq data was processed with the R package Seurat (version 3.1.5) (15). The cells were removed that had either fewer than 301 expressed genes or over 30% UMIs originating from mitochondrial. UMI counts were normalized and were transformed to the log-transformed. Three thousand highly variable Genes (HVGs) of each data were identified. Integration of single-cell data was performed to correct batch effect. Principal component analysis (PCA) and uniform manifold approximation and projection (UMAP) dimension reduction were performed with top 14 principal components. The Louvian modularity optimization algorithm was applied to iteratively group cells together into clusters. Cell clusters were annotated to known biological cell types using canonical cell marker genes.

### Differential gene expression analysis

To identify genes differentially expressed in COPD compared to control in each cell type, MAST(1.14.0) (16) was used to perform zero-inflated regression analysis by fitting a linear mixed model. We controlled both technical variation and individual variation by using a two-part hurdle model. The following model was fit with MAST:

zlm(∼ COPD+ age + (1|individual) + smoking + percent.mt + nCount_RNA, sca, method=’glmer’, ebayes=FALSE)

Where nCount_RNA was the total number of molecules detected within a cell and percent.mt was mitochondria fraction. To identify genes differentially expressed due to the disease effect, likelihood ratio test was performed by comparing the model with and without the COPD variable. Multiple hypothesis testing correction was performed with Bonferroni and Holm corrections. Genes with FDR < 0.05 were selected as differentially expressed genes (DEGs). Similar approach was applied to age and smoking variable.

### Down-sampling analysis to identify condition-relevant cell types

To calculate the number of COPD-associated DEGs across cell types normalized by number of cells in each cell cluster, we randomly drew 700 cells from each cell cluster before performing differential gene expression analysis. This analysis was repeated 10 times and the number of DEGs for each cell cluster were presented as boxplots. Age-associated DEGs and smoking-associated DEGs were performed as above.

### Subclustering and identification of markers of subcluster

To identify subclusters within innate immune cells, we reanalyzed cells belonging to monocytes and dendritic cells with a resolution of 0.2 separately. Subclusters were annotated to known biological cell types using canonical cell marker genes or to cell types of published literature with their cell marker genes.

To identify marker genes for each of subclusters within monocytes, we contrasted cells from that subcluster to other cells in monocytes using a Wilcoxon Rank Sum test. Additionally, marker genes were required to have the highest mean expression in that subcluster. Marker genes for each of subclusters within dendritic cells were analyzed like monocytes.

### Enrichment analysis of gene sets

Gene Ontology Gene enrichment was performed using the R package clusterProfiler (version 3.14.3) (17). Enrichment in GO Biological process between DEGs of COPD of each cell cluster were analyzed using the compareCluster. Significantly enriched GO terms were simplified using simplify remove highly similar terms (cutoff = 0.7) by retaining the most significant representative term. Enrichment in GO Biological process on DEGs of age and smoking were analyzed as above.

Enrichment analysis using custom genes sets was performed using the appropriate background lists (genes detected >10 % of cells in each cell cluster). Enrichment of COPD-associated DEGs at COPD GWAS risk genes (18) was performed using Fisher’s exact test. Similarly, enrichment of subcluster’s markers at COPD-associated DEGs or age-associated DEGs or smoking-associated DEGs were analyzed as above.

### Trajectory analysis

To identify lineage trajectories, we used Slingshot (version 1.4.0) (19). The cluster representing alveolar type 2 cells and club cells were chosen as the root node separately for calculation of lineages and pseudotime with PCA-based dimension reduction was performed with 3000 HVGs.

After running slingshot, we used the tradeSeq package (version 1.4.0) (20) to find genes dynamically expressed during cell differentiation. For each HVG, we fit a general additive model (GAM) using a negative binomial noise distribution to model the relationships between gene expression and pseudotime and tested for significant associations between expression and pseudotime using the associationTest. We picked out the top genes based on wald statistics and visualized their expression over developmental time with a heatmap.

### Cell-cell communication analysis

Cell-cell communication analysis was conducted with the CellPhoneDB software (version 2.0) (21). Only receptors and ligands expressed in >10% of cells from COPD or elderly control or young control were further evaluated, while a cell-cell communication was considered nonexistent if the ligand or the receptor was unmeasurable. Averaged expression of each ligand-receptor pair was analyzed between various cell clusters, and only those with P value < 0.05 were used for the prediction of cell-cell communication between any two cell clusters.

## Results

### scRNA-seq and cell typing of COPD and non-COPD lung tissues

Nine patients with untreated, primary, non-metastatic NSCLS of adenocarcinoma subtype that underwent lung lobe resection with curative intent were divided into 3 groups according to lung function and age, namely COPD group, control old group, and control young group. Among them, 4 patients were active smokers and 5 others were non-smokers (Supplementary Table 1). After resection, one non-malignant lung tissue sample from a distal region within the resected lobe was obtained, rapidly digested to a single-cell suspension, and analyzed via scRNA-seq. After data processing, we classified cells into 15 groups of cell types, which included 14 known cell types (T cells, B cells, neutrophils, natural killer cells (NKs), monocytes, mast cells, macrophages, dendritic cells (DCs), stromal cells, endothelial cells, club cells, ciliated cells, alveolar type 1 cells, and alveolar type 2 cells) and a distinct unknown cluster highly positive for cell proliferation markers which was labeled as proliferating cells (Fig. 1A).

**Figure 1.**
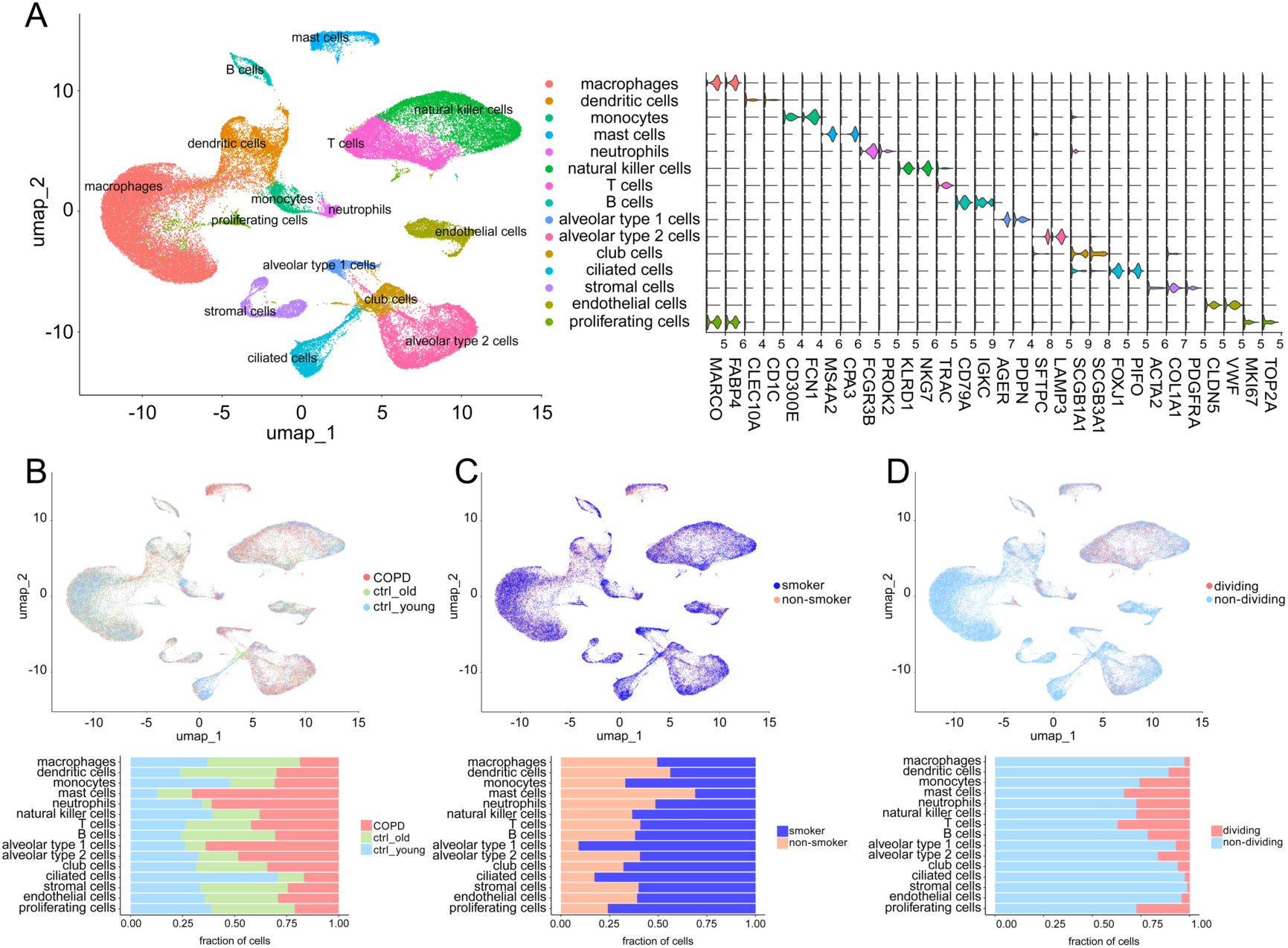
Overview of the scRNA-seq dataset from COPD and non-COPD lung tissue samples. (**A**) Clustering of the 65,000 cells, with each cell clolorcoded for the associated cell type (left). The violin plots displayed the expression distributions of known marker genes according to which cell types were annotated (right). (**B**) tSNE plot from Fig. 1A colored by group label (up). The fractions of all cell types originating from each group were showed in the boxplot (bottom). (**C**) tSNE plot from Fig. 1A colored by the smoking status of the patients (up). The fractions of all cell types originating from smoking and non-smoking patients were showed in the boxplot (bottom). (**D**) tSNE plot from Fig. 1A colored by cell dividing state (up). The fractions of all cell types originating from dividing and non-dividing cells were showed in the boxplot (bottom).

Each cell type was then subdivided according to group assignment (group, Fig. 1B), smoking status of patients (smoking, Fig. 1C), and cell dividing state (cell cycle, Fig. 1D), and the percentages of all subtypes were calculated. Taking age into consideration, we noticed that compared with control young group, only dendritic cells, monocytes, and ciliated cells in the COPD group displayed consistent trends of change with those in control old group (Fig. 1B), indicating that the pathogenesis of COPD might not simply be accelerated and/or aberrant lung aging. When analyzing the effect of smoking on each cell type, we observed that as expected, the abundance of most structural cells, especially alveolar type 1 cells, ciliated cells, and club cells were reduced in the lung tissues from smokers, compared with those in lung tissues from non-smokers. As for immune cells, only mast cells and dendritic cells showed significant increase in quantities in the lung tissues from smokers (Fig. 1C). Cell types predicted to be capable to divide were as expected, such as most immune cells usually with relatively high turnover rates, along with several kinds of structural cells with well-established potential of self-renewal and differentiation, including alveolar type 2 cells (AT2s) and club cells (Fig. 1D).

### Cell type-specific gene expression changes in COPD lung tissues

Normalized numbers of differentially expressed genes (DEGs) under three different pathological conditions showed great distinctions within several cell types. Likewise, numbers of DEGs under certain pathological condition also significantly differed from each other among some cell types (Fig. 1A). We performed Gene Ontology (GO) analysis of aging-(age, old vs. young), COPD-(COPD, COPD vs. ctrl), and smoking-(smoking, smoking vs. non-smoking) associated DEGs, respectively (Fig. 2A). As expected, many GO terms related to classical hallmarks of aging, including loss of proteostasis, genomic instability, epigenetic alterations, and mitochondrial dysfunction, were found among aging-associated DEGs, reflecting the general status of lung aging from the elderly subjects. Similarly, pathways of T cell activation, response to reactive oxygen species (ROS), response to oxidative stress (OS), and response to endoplasmic reticulum stress (ERS) were enriched in smoking-associated DEGs, suggesting the existence of cellular stress under the condition of smoking. Meanwhile, regulation of response to cytokine stimulus, regulation of cytokine production, and regulation of innate immune response were among the top dysregulated pathways in COPD-associated DEGs, implying the involvement of cytokine and innate immune system in the development of COPD. In searching of the cellular origin of COPD-specific DEGs, we divided all the clusters except for proliferating cells into immune and non-immune cells. Among immune cells, the top COPD-associated DEGs were from monocytes; among non-immune cells, they were from epithelial and endothelial cells (Fig. 3B). Additionally, the intersection between COPD-specific DEGs and the COPD risk genes from genome-wide association study (GWAS) (18) showed that the GWAS COPD risk genes were mostly overrepresented in monocytes, followed by alveolar type 2 cells (AT2) and endothelial cells (Fig. 2C). Differential expression burden analysis showed that monocytes were preferentially affected in COPD, while club cells and macrophages were the most relevant types under aging and smoking, respectively. Intriguingly, we also noticed smoking-associated changed were the strongest in endothelial cells among non-immune cells (Fig. 2D). Together, these results highlighted the cell type-specific changes under different conditions, particularly monocytes in COPD lungs, with the enrichment in cytokines and innate immune system.

**Figure 2.**
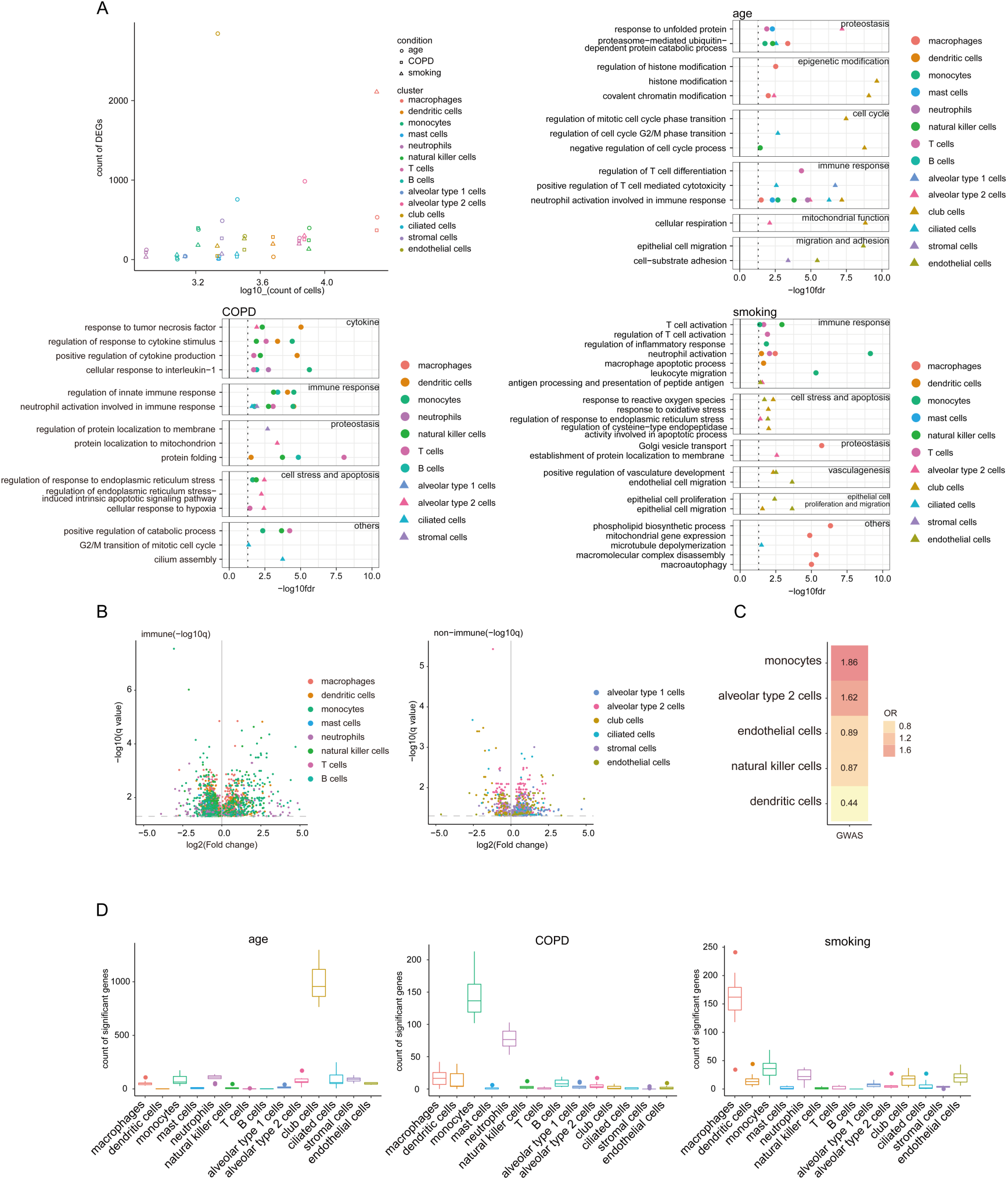
Cell type-specific gene expression changes in COPD lung tissues. (**A**) The numbers of DEGs in old lung tissues vs. young lung tissues (age, circle), COPD lung tissues vs. control lung tissues (COPD, square), and lung tissues from smokers vs. lung tissues from non-smokers (smoking, triangle) of each cell type were normalized with cell counts (top left). GO analysis of age-(top right), COPD-(bottom left), and smoking-(bottom right) associated DEGs shows the enrichment of biological pathways under each condition. (**B**) Volcano plots for COPD-associated DEGs expressed in immune cells (left) and non-immune cells (right). (**C**) Overlap between cell type-specific COPD-associated DEGs and GWAS COPD risk genes. (**D**) Burden analysis was carried out by comparing aging-associated DEGs (left), COPD-associated DEGs (middle), and smoking-associated DEGs (right) across all cell types.

**Figure 3.**
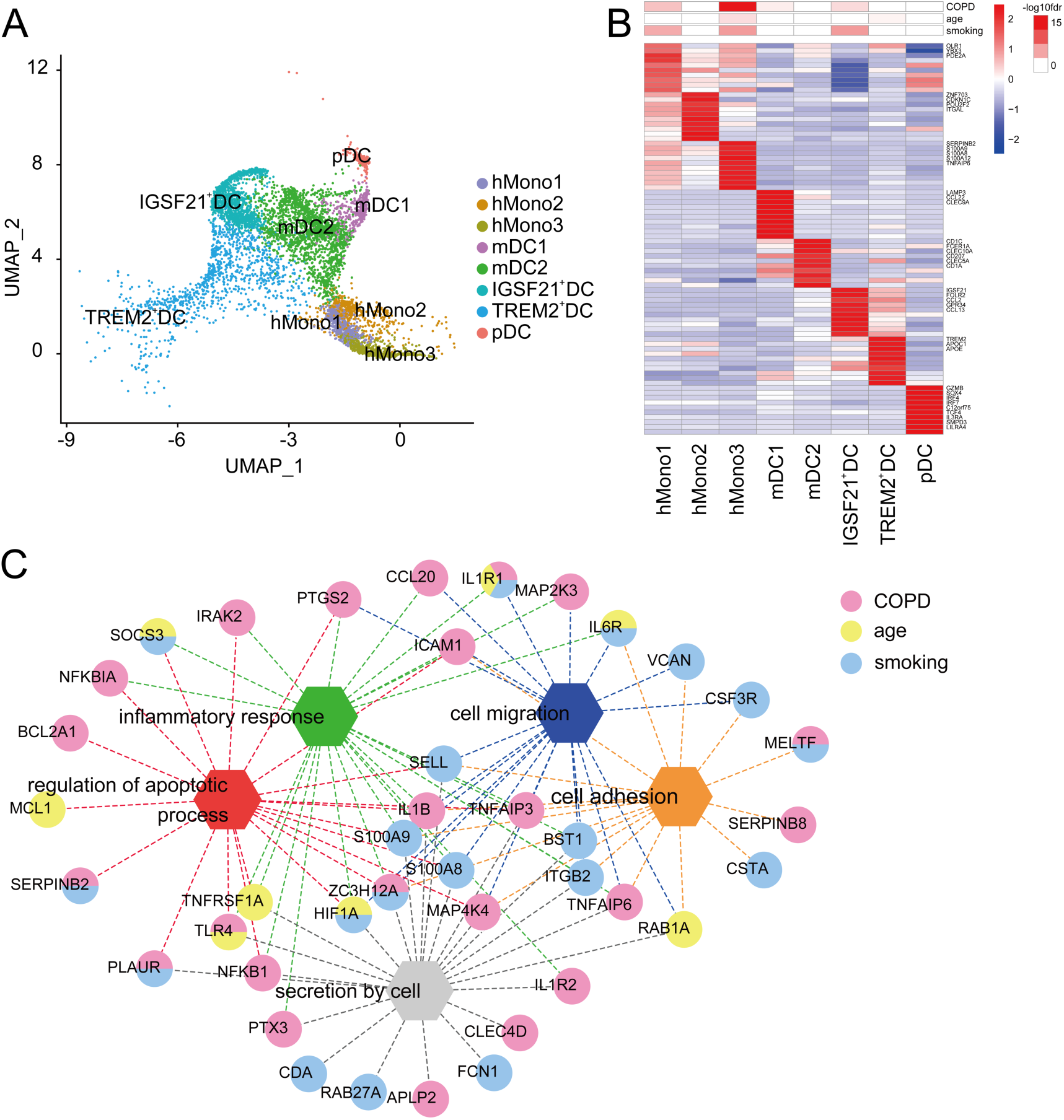
Abnormal activation of monocytes in COPD lung tissues. (**A**) SPRING plots of monocyte and DC subsets. Cells were colored by inferred cell type. (**B**) Genes enriched between monocyte and DC subsets (bottom). Distribution of DEGs under three conditions (COPD, age, and smoking) in each subset of monocyte and DC (top). (**C**) Functional interaction network generated by the STRING database for the hMono3 marker genes overlapped with monocytes DEGs under different conditions (COPD, age, and smoking), with nodes representing genes, edges representing interactions, and colors representing conditions. Disconnected genes were not shown.

### Abnormal activation of monocytes in COPD lung tissues

The heterogeneity of mononuclear phagocyte system (MPS) has long been proposed and was recently verified by various studies using single-cell sequencing (22). Here, we defined 8 transcriptional states of monocytes and DCs in reference to annotation in 2 recent studies (23, 24), including 3 subsets of monocytes (hMono1-3) and 5 subsets of DCs (mDC1-2, IGSF21^+^DC, TREM2^+^DC, and pDC) (Fig. 3A), each of which showed distinct gene expression profiles (Fig. 3B). In fact, hMono1, hMono2, and hMono3 correspond well to classical (CD14^+^CD16^-^), non-classical (CD14^dim^CD16^+^), and intermediate monocytes (CD14^+^CD16^+^) described in humans, respectively (25). Similar to the canonical categorization of DCs into three independent developmental lineages including conventional DCs (cDCs), plasmacytoid DCs (pDCs), and monocyte-derived DCs (moDCs) (26, 27), the mDC1 in our data expressed cDCs-associated genes like LAMP3 (28), and the pDC here expressed pDCs markers, such as IRF4 (28). Besides, we also identified 3 non-canonical DC clusters according to their distinctive marker genes (Fig. 3B). Among monocyte subtypes, we observed enrichment of COPD-associated DEGs in hMono3 (Fig. 3B), suggesting hMono3 play a critical role in COPD pathogenesis. Consequently, to gain better insights into the contribution of hMono3 in COPD development, we created a StringDB network based hMono3-specific DEGs in all three conditions (COPD, aging, and smoking), which revealed regulation of apoptosis process as the most relevant biological pathway among hMono3-related COPD-associated DEGs, together with cell migration and secretion (Fig. 3C). These findings highlight the abnormal activation of monocytes and subsequent disruption in regulating apoptosis process in COPD lungs, stressing its potential killing effect on other cells in the development of COPD.

### Apoptosis of alveolar epithelial cells in COPD lung tissues and exhaustion of club cells stemness under aging conditions

Increased alveolar wall destruction in COPD lungs has long been attributed to the reduction in both quantity and regenerative capacity of airway stem/progenitor cells including AT2s and club cells (29). As expected, slingshot trajectory of AT1s and AT2s in our data illustrated transition from AT2s to AT1s (Fig. 4A), supporting the capability of AT2s to self-renew and differentiate into AT1s (29), which was reconfirmed by the enrichment of AT2s in earlier trajectory along the pseudo-timeline compared with AT1s (Supplementary Fig. S1). Notably, AT2s in COPD lungs appeared “less differentiated” in general, compared with those in control lungs (Fig. 4B), suggesting the suppression of AT2 to AT1 differentiation in COPD lungs. We identified dynamically altered gene in expression during transition, which were potentially key regulators for AT2 differentiation (Fig. 4C), such as two classical apoptosis-associated genes, namely HES1 (30) and KLF4 (31) (Fig. 4D). Intriguingly, both HES1 and KLF4 significantly increased in AT2 from COPD lungs compared with controls (Fig. 4E), indicating the contribution of AT2 apoptosis to alveolar epithelium destruction in COPD lungs.

**Figure 4.**
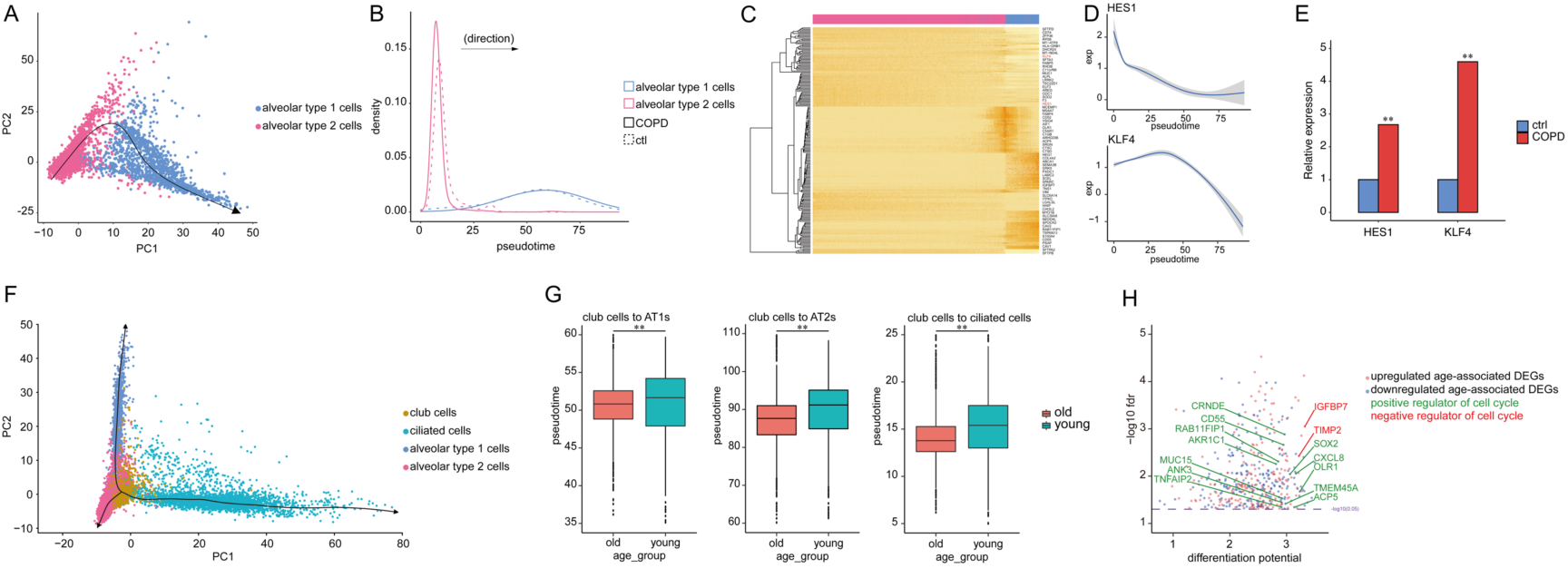
Apoptosis of alveolar epithelial cells in COPD lung tissues and exhaustion of club cells stemness under aging conditions. (**A**) A trajectory of AT2-to-AT1 differentiation. (**B**) Pseudotemporal cell density dynamics of AT1s (blue) and AT2s (red) in control (dotted line) and COPD (full line) lungs. (**C**) Heatmaps displaying the scaled expression profile for each gene (rows) with cells (columns) ordered according to their differentiation positions. (**D**) Dynamics of gene expression of HES1 and KLF4 along the pseudo-timeline. (**E**) Relative gene expression levels of HES1 and KLF4 in control and COPD AT2s (** p < 0.05 vs. ctrl). (**F**) A trajectory of airway epithelial cells differentiation including club cells (yellow), ciliated cells (green), AT1s (purple), and AT2s (red). (**G**) Pseudotime of club cells during club cells-to-AT1s (left), club cells-to-AT2s (middle), and club cells-to-ciliated cells (right) differentiation in both young and old lungs (** p < 0.05 vs. young). (**H**) Scatter plot for age-associated DEGs with significant differentiation potential in club cells.

As shown above, club cells were preferentially affected as ages (Fig. 2D). Given that club cells are a bronchoalveolar epithelial cells with multiple functions including regeneration, which is also of great importance to alveolar maintenance (32, 33), we hypothesized that aged club cells may be restrained from differentiation, which facilitates alveolar wall destruction during COPD development. To test this hypothesis, we first inferred pseudo-temporal ordering of club cells, ciliated cells, AT1s, and AT2s differentiation trajectories in a two-dimensional t-SNE map (Fig. 4F), confirming the capability of club cells to differentiate to the other three kinds of cell types. Note that club cells in aged lungs were more enriched in earlier trajectory than that in young lungs in all the three differentiation modes (Fig. 4G), implying either the impaired differentiation ability or the dysregulated self-renewal of club cells. Since aging-associated DEGs were enriched in biological pathways of cell cycle (Fig. 2A), we therefore took a deep look into the expression of cell cycle-related genes to identify potential genes responsible for suppression of club cell differentiation. We found that several pro-proliferative DEGs (including OLR1 (34), CXCL8 (35), ACP5 (36), SOX2 (37), TMEM45A (38), CRNDE (39), CD55 (40), and HSPA2 (41) were downregulated, while some anti-proliferative ones (including IGFBP7 (42) and TIMP2 (43)) were elevated in aged club cells compared to those in young club cells, respectively (Fig. 4H), which strongly implied the dysregulation of aged club cells in terms of self-renewal. In sum, our findings demonstrated the malfunction of both AT2s and club cells as stem/progenitor cells in lung epithelium, and revealed the elevated activity of apoptosis in AT2s accounting for alveolar wall destruction in COPD lungs and the decline of aged club cell in self-renewal contributing to the development of COPD.

### Enhanced intercellular communication in COPD lung tissues and pro-inflammatory macrophage-endothelial cell interaction in the context of smoking

Altered intercellular communication has long been considered as a critical hallmark of lung aging that plays a role in COPD (9); however, there fell short of a close look of the interaction between various cell types within COPD lung tissues. We therefore sought to identify abnormal intercellular communication between various cell types at single-cell resolution in COPD lungs. A panoramic view of the interactions of the 15 clusters in the lung tissues from young and old control groups as well as COPD group clearly showed that cellular interactions between innate immune cells (especially monocytes) and non-immune structural cells (including AT2s, endothelial cells, club cells, AT1s, and stromal cells) tended to enhance with age, and was even stronger in COPD lung tissues (Fig. 5A). Since most of intercellular communications depend on interactions between specific ligands and receptors expressed by communicating cells, we subsequently investigated the ligand-receptor pairs between several typical cell types showing enhanced intercellular communications in COPD lungs (Fig. 5B). Notably, some ligand-receptor pairs, such as IL6-IL6 receptor pair between monocyte and AT2 that was both up-regulated in expression level and increased in the intensity of interaction, may not only contribute to the enhanced intercellular communication between monocytes and AT2s in COPD lungs, but also account for the damaging effect of monocytes on AT2s in COPD development.

**Figure 5.**
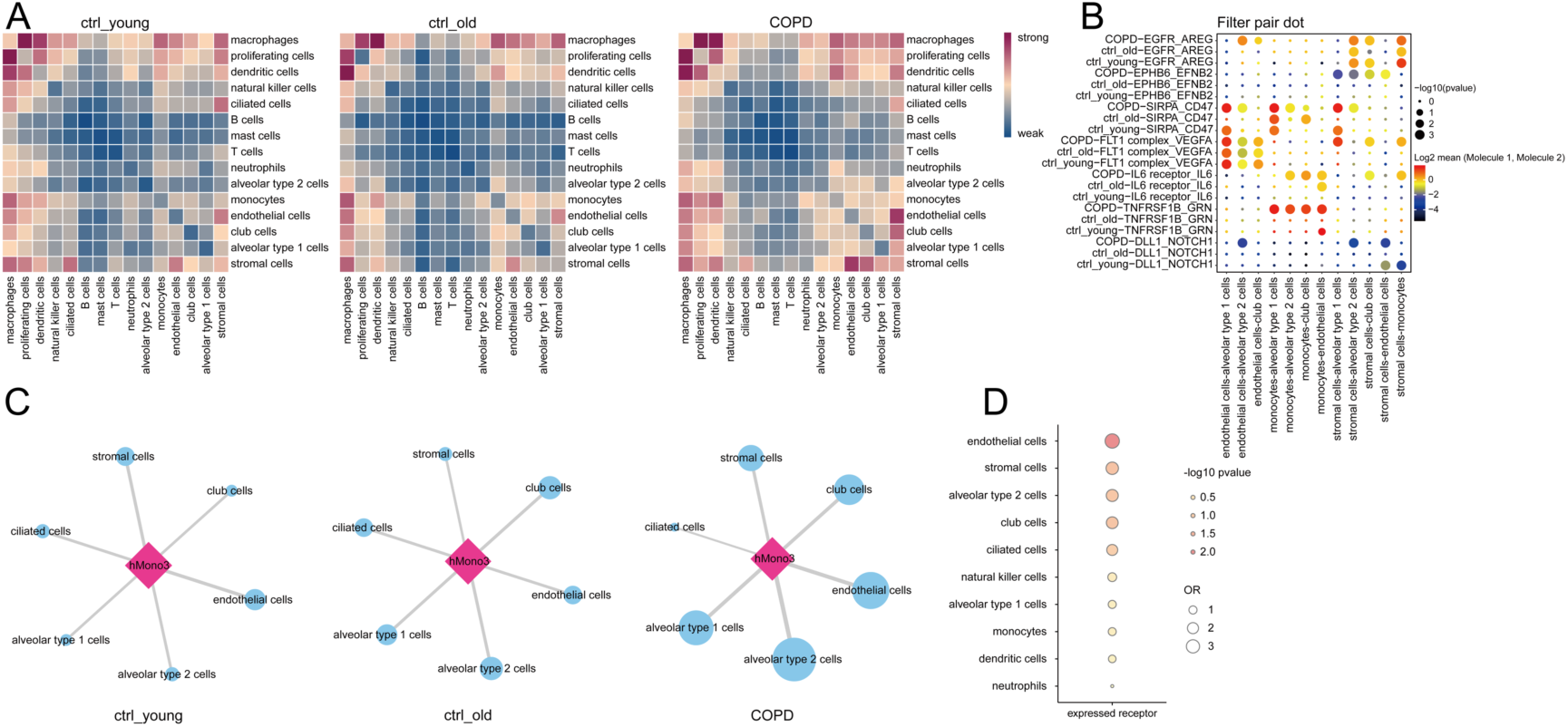
Enhanced intercellular communication in COPD lung tissues and pro-inflammatory macrophage-endothelial cell interaction in the context of smoking. (**A**) Heatmap showing the intensity of interactions between all cell types in lungs from young controls (ctrl_young, left), old controls (ctrl_old, middle), and COPD patients (COPD, right). (**B**) Representative ligand– receptor pairs which may account for the interactions between certain cell types in lungs from young controls (ctrl_young), old controls (ctrl_old), and COPD patients (COPD). P values are indicated by circle size; scale is shown beside the plot. The means of the average expression level of interacting molecule 1 in cluster 1 and interacting molecule 2 in cluster 2 are indicated by color. (**C**) The intensity of interactions between hMono3s and other structural cells including AT1s, AT2s, endothelial cells, club cells, stromal cells, and ciliated cells in lung tissues from both young and old controls as well as COPD patients. (**D**) The intensity of interactions between macrophages and other cell types in smoking lungs. P values are indicated by color. Odds ratio (OR) is indicated by circle size; scale is shown beside the plot.

Next, we sought to infer the potential consequence of abnormal activity of the hMono3 by examining their interaction with and other structural cells in COPD lungs. As can be seen from Fig. 5C, under the conditions of COPD, the interactions between hMono3s and AT2s as well as AT1s were significantly increased, followed by those between hMono3s and endothelial cells as well as club cells, when compared with lungs from both young and old normal controls, which again strongly suggested the potential abnormal damaging effect of abnormally activated monocytes on alveolar epithelium during the development of COPD.

As macrophages and endothelial cells turned out to have the most of smoking-associated changes in immune and non-immune cells, respectively (Fig. 2D), we hypothesized that macrophages could actively interact with endothelial cells under smoking conditions. As expected, cell-cell interaction analysis conducted via CellPhoneDB showed macrophages have the highest interaction with endothelial cells (Fig. 5D), with the predicted ligand-receptor pairs playing the leading role in such intensified macrophage-endothelial cell communication (Supplementary Table S2), which is very likely to promote the accessibility of alveolar epithelium to aberrantly activated monocytes via increasing the permeability of pulmonary vascular endothelium, mediating the contribution of smoking to COPD pathogenesis. Together, these findings highlight the enhanced intercellular communication in COPD lungs and uncover the active interaction between macrophages and endothelial cells by smoking, which is very likely to promote the development of COPD.

## Discussion

The present study, as far as we know, is the first to perform in-depth analysis of scRNA-seq data from COPD lung tissues, while taking both aging and smoking into account. Our results showed the enhanced intercellular communication in COPD lung tissues underpinned the crosstalk between abnormally activated monocytes and pulmonary structural cells, especially AT2s, which aggregately indues AT2 apoptosis and the subsequent alveolar epithelium injury. Additionally, trajectory analysis demonstrated club cells have the capacity to repair the alveolar epithelium damage via differentiating into AT1s and AT2s, which were suppressed as ages. Furthermore, as the first- and second-ranked smoking relevant cell types indicated by cell burden analysis, macrophages and endothelial cells showed enhanced interaction with each other in smoking lung tissues, which is characterized by bidirectional pro-inflammatory effects, leading to the progression of the chronic inflammation responsible for the damage of alveolar epithelium during COPD development. Our findings uncover the highly cell type-specific changes in different pathological settings (COPD, aging, and smoking) and the internal connections of them from the perspective of intercellular communication at a single-cell resolution, responsible for the potential mechanisms by which aging and smoking facilitate COPD pathogenesis.

In the past, adaptive immune system has been extensively studied in investigations of COPD pathogenesis due to the conventional view that the adaptive immune response is crucial for the type of long-term inflammation that is required to drive chronic respiratory disease such as COPD. Although mounting evidences are emerging that the participation of innate immune system in chronic respiratory disease are at least equally important (44–46), monocytes have long been unappreciated mainly due to its plasticity to both DCs and macrophages. Moreover, the heterogeneity of this cellular population further increased the difficulty of research. Travaglini et al. identified three major subclusters of monocytes in their recently published molecular cell atlas of the human lung from scRNA-seq, corresponding to classical, non-classical, and intermediate monocytes, respectively (23). Similarly, we also annotated the monocytes in our data as hMono1 (classical monocyte), hMono2 (non-classical monocyte), and hMono3 (intermediate monocyte). Enrichment of COPD-associated DEGs in hMono3s among the three monocyte subclusters indicated its vital in COPD pathogenesis. In accord with the role for intermediate monocytes in antigen presentation, cytokine secretion, apoptosis regulation, and differentiation (25), we also found the enrichment of regulation of apoptotic process and secretion by cell in the COPD-associated DEGs of hMono3s (such as TLR4 (47) and IL1B (48)), which raised the possibility that hMono3s may serve as the aberrantly activated subcluster of monocytes in the COPD lungs and have a killing effect on other cell types through abnormally secreting cytokines. Meanwhile, AT2s in COPD lungs showed not only the inhibited differentiation to AT1s, but also increased expression of several positive apoptotic regulators in its COPD-associated DEGs, such as HES1 (30) and KLF4 (31). Thus, AT2s were the potential killing target of hMono3s. Expectedly, CellPhoneDB analysis demonstrated enhancement of interaction between monocytes and AT2s in COPD lung tissues with several ligand-receptor pairs whose expressions and interactions significantly increased, including CCL3L1-DPP4. Interestingly, as a potent CCR5 binding chemokine, CCL3L1 can be cleaved by DPP4 into a truncated form, with an enhanced chemotactic activity (49). Given that hMono3 is the only subset of monocyte expressing CCR5 (25), such chemokine-protease interaction between CCL3L1 and DPP4 is very likely to exert a positive feedback recruiting more hMono3s to alveolar epithelium in COPD lung tissues. Moreover, we also noticed that LGALS9 encoding pro-apoptotic galectin-9 (50) was among the monocyte-derived ligands interacting with receptors from alveolar epithelial cells. Further experiments are needed to validate the existence as well as the exact roles of these interactions; however, our results highlight the biological significance of monocytes, essential but previously unappreciated component of innate immune system, in COPD pathogenesis, greatly expanding the current pathological model of COPD centered on macrophages.

Previous studies have uncovered more than one kind of progenitor cells participating in alveolar epithelium regeneration, including type 2 alveolar epithelial cells (AT2s) (51, 52) and CCSP^+^ club cells (also known as clara cells) (53). Although the pluripotency of club cells has been observed in various murine models (54, 55), it has not been fully verified in human airway epithelium. Here, to our knowledge, we are the first to reveal the multipotential differentiation of club cells in human lung by trajectory analysis based on scRNA-seq data, which require experimental validation. Zuo et al. identified novel roles for human club cells in host defense, xenobiotic metabolism, antiprotease, physical barrier function, monogenic lung disorders, and receptors for human viruses and provided a batch of functional genes expressed by human club cells (56). We noticed that these genes considerably overlapped with the aging-associated DEGs of club cells in our data, suggesting the degeneration in club cell physiology that is consistent with stem cell exhaustion, one of the classical hallmarks of aging (9, 57). Consistently, we found that IGFBP7 (42) and TIMP2 (43), which negatively regulate cell proliferation increased, while genes promoting cell growth including OLR1 (34), CXCL8 (35), ACP5 (36), SOX2 (37), TMEM45A (38), CRNDE (39), CD55 (40), and HSPA2 (41) decreased in aged club cells, revealing the dysregulation of aged club cells in proliferation. Given the essential role of club cell stemness in airway epithelium repair, such dysregulation is very likely to aggravate the alveolar epithelium injury in COPD lungs as ages.

In our current study, macrophages were defined as the most smoking-associated cell type followed by endothelial cells in down-sampling analysis, highlighting the vital role of macrophages under the condition of smoking. Interestingly, the enhanced cellular interaction between macrophages and endothelial cells is supported by cell-cell communication analysis. Thus, such interaction in the context of smoking may be harmful to the integrity of vascular endothelium and even survival of endothelial cells, since several ligand-receptor pairs responsible for the enhanced macrophage-endothelial cell interaction in smoking lung tissues were thought to mediate the detrimental role of macrophages in vascular homeostasis. For example, PDGFC encoding PDGF-CC which is a recently discovered high-affinity ligand of VEGFR2 (58), was significantly up-regulated in macrophages from smoking lung tissues compared to control lung tissues. In our data, PDGF-CC is very likely to increase vascular permeability via interacting and activating VEGFR2 on the cell membrane of endothelial cells which may lead to the dissociation and internalization of the complex (59, 60). Additionally, the Sema3E (Semaphorin 3E)-PlexinD1 interaction with suppressive effect on endothelial cell proliferation and angiogenic capacity in brain (61) was also shown to be enhanced between macrophages and endothelial cells in smoking lung tissues by CellPhoneDB analysis. Moreover, the Semaphorin 3E secreted by macrophages was reported to inhibit macrophage migration, therefor promoting macrophage retention and chronic inflammation in atherosclerosis (62), which suggests the amplification of local inflammation by Sema3E-PlexinD1 interaction in an autocrine manner. Further experiments are needed to elucidate the exact mechanisms and effects of the communication between macrophages and endothelial cells in lung tissues enhanced by smoking. In sum, our results suggested macrophage still play an important role in the development of COPD, however, in a smoking-dependent manner.

There are certainly some limits of our current study, including 1) sample collection based on non-tumorous adjacent lung tissues, 2) limited number of subjects involved, and 3) the age bias of patients with COPD. Although bringing new insights into the pathological mechanisms of COPD, our findings need to be fully verified by future in vivo experiments.

In summary, via analyzing the scRNA-seq data of the lung tissues from COPD patients, young control, and elderly control individuals, we identified monocytes, club cells, and macrophages as the most COPD-, aging-, and smoking-associated cell types. Advanced bioinformatic analyses further revealed the cell-type specific changes under these different pathologic conditions, implying the possible mechanisms by which aging and smoking promote COPD development through club cell stemness exhaustion and pro-inflammatory macrophage-endothelial cell interaction, respectively. Our research provides a new paradigm for the study of the COPD pathogenesis, makes an important supplement to the existing pathological mechanisms, and is expected to bring new strategies and targets for the prevention and treatment of COPD.

## Author contributions

The study was designed by Q.H., M.L., and J.W.. Q.H., S.S., Y.C., and S.W. collected the samples. Q.H., S.S., and Y.C. performed the experiments. M.L., J.W., Y.W., and Q.H. did the bioinformatic analyses. Q.H., M.L., and J.W. wrote the manuscript. Additional expertise was contributed by W.X. and B.C..

## Acknowledgements

We are grateful to Xiaosu Li, a freelance translator in Beijing, China, for editorial assistance.

