## Supplementary Information for "Single-cell transcriptomics reveal cell type-specific molecular changes and altered intercellular communications in chronic obstructive pulmonary disease"

**Supplementary Table S1.** Characteristics of the 9 patients included in this study.

| Patient | Gender | Age | COPD (GOLD) | Pathologic type | Affected lobe | Smoking status |
|---|---|---|---|---|---|---|
| 1 | Male | 73 | No | pneumocyte hyperplasia | Right middle | Active |
| 2 | Male | 71 | No | Adenocarcinoma | Right upper | Active |
| 3 | Male | 63 | Yes (II) | Adenocarcinoma | Left down | Active |
| 4 | Male | 28 | No | Adenocarcinoma | Left upper | Non-smoker |
| 5 | Male | 75 | No | Adenocarcinoma | Left down | Non-smoker |
| 6 | Male | 73 | Yes (II) | Adenocarcinoma | Right middle | Non-smoker |
| 7 | Male | 50 | Yes (III) | Adenocarcinoma | Left down | Active |
| 8 | Male | 35 | No | Adenocarcinoma | Right down | Non-smoker |
| 9 | Male | 27 | No | Adenocarcinoma | Right middle | Non-smoker |

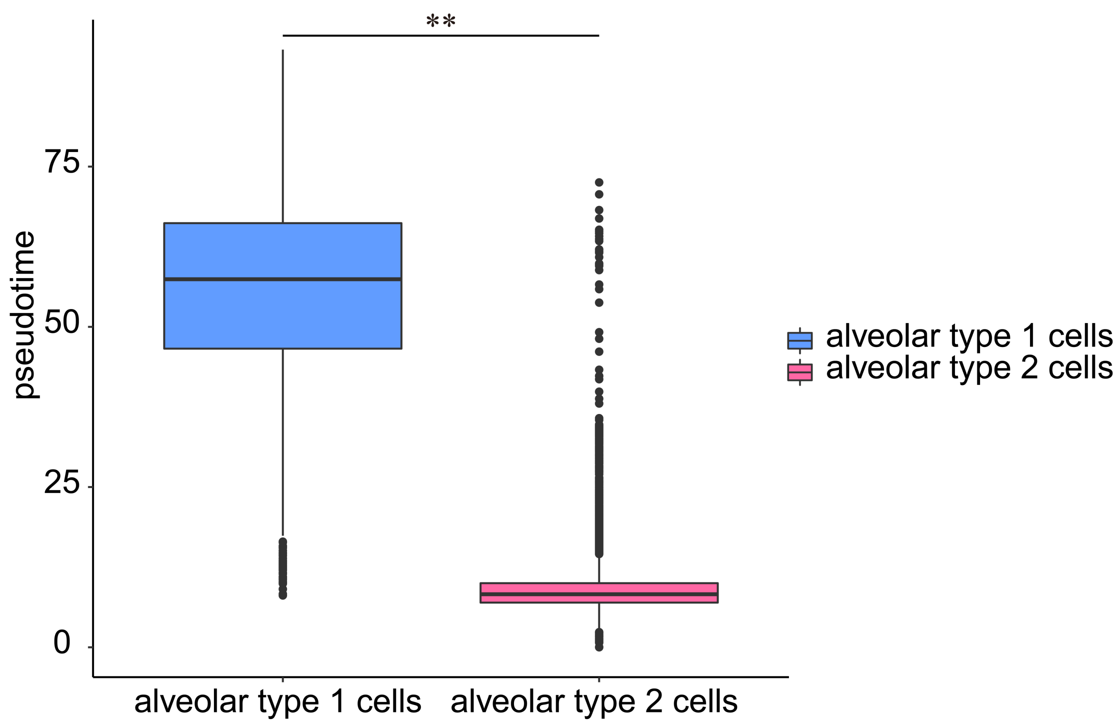


**Supplementary Figure S1.**

Pseudotime of AT1s and AT2s during AT2s-to-AT1s differentiation in lung tissues (** $p < 0.05$ vs. alveolar type 1 cells).

**Supplementary Table S2.** The predicted ligand-receptor pairs that are most responsible for the intercellular communication between macrophages and endothelial cells in lung tissues intensified by smoking.

| **Ligand (macrophage)** | **Receptor (endothelial cell)** | **pvalue (ligand)** | **log2fc (ligand)** | **receptor expression percent** |
|---|---|---|---|---|
| PDGFC | FLT4 | 3.59E-06 | 1.60959395 | 0.263008515 |
| PDGFC | KDR | 3.59E-06 | 1.60959395 | 0.306212551 |
| SEMA3E | PLXND1 | 3.16E-06 | 0.609719309 | 0.240618102 |
| TGFB1 | ACVRL1 | 0.044363521 | 0.063956487 | 0.452538631 |
| TGFB1 | ENG | 0.044363521 | 0.063956487 | 0.281929991 |
| TGFB1 | TGFBR2 | 0.044363521 | 0.063956487 | 0.620309051 |
| TGFB1 | TGFBR3 | 0.044363521 | 0.063956487 | 0.443077893 |
